# Reliability assessment of temporal discounting measures in virtual reality environments

**DOI:** 10.1101/2020.08.07.237826

**Authors:** Luca R. Bruder, Lisa Scharer, Jan Peters

## Abstract

In recent years the emergence of high-performance virtual reality (VR) technology has opened up new possibilities for the examination of context effects in psychological studies. The opportunity to create ecologically valid stimulation in a highly controlled lab environment is especially relevant for studies of psychiatric disorders, where it can be problematic to confront participants with certain stimuli in real life. However, before VR can be confidently applied widely it is important to establish that commonly used behavioral tasks generate reliable data within a VR surrounding. One field of research that could benefit greatly from VR-applications are studies assessing the reactivity to addiction related cues (cue-reactivity) in participants suffering from gambling disorder. Here we tested the reliability of a commonly used temporal discounting task in a novel VR set-up designed for the concurrent assessment of behavioral and psychophysiological cue-reactivity in gambling disorder. On two days, thirty-four healthy non-gambling participants explored two rich and navigable VR-environments (neutral: café vs. gambling-related: casino and sports-betting facility), while their electrodermal activity was measured using remote sensors. In addition, participants completed the temporal discounting task implemented in each VR environment. On a third day, participants performed the task in a standard lab testing context. We then used comprehensive computational modeling using both standard softmax and drift diffusion model (DDM) choice rules to assess the reliability of discounting model parameters assessed in VR. Test-retest reliability estimates were good to excellent for the discount rate log(k), whereas they were poor to moderate for additional DDM parameters. Differences in model parameters between standard lab testing and VR, reflecting reactivity to the different environments, were mostly numerically small and of inconclusive directionality. Finally, while exposure to VR generally increased tonic skin conductance, this effect was not modulated by the neutral vs. gambling-related VR-environment. Taken together this proof-of- concept study in non-gambling participants demonstrates that temporal discounting measures obtained in VR are reliable, suggesting that VR is a promising tool for applications in computational psychiatry, including studies on cue-reactivity in addiction.

## Introduction

Recent research has exploited the development of high-performance virtual reality (VR) technology to increase the ecological validity of stimuli presented in studies of cue-exposure^[1–3]^, counterconditioning^[4]^, equilibrium training^[5]^, social gazing^[6]^ and gambling behavior in healthy control participants^[7]^. Furthermore, it has been shown to increase immersion and arousal during gambling games^[8]^. However, before VR can be widely applied with confidence it is important to establish that commonly applied behavioral tasks still yield reliable data in a VR context. Research focusing on psychiatric disorders, where one goal is to create reliable diagnostic markers based behavioral tasks and model-based computational approaches, would benefit from behavioral tasks that produce reliable parameters on a single participant level in VR.

A core characteristic of many psychiatric and neurological disorders is a detrimental change in decision-making processes. This is especially evident in addiction-related disorders such as substance abuse^[9–11]^ or gambling disorder^[12–14]^. One approach to study such changes in decision making is computational psychiatry^[15]^, which employs theoretically grounded mathematical models to examine cognitive performance in relation to psychiatric disorders. Such a model-based approach allows for a better quantification of the underlying latent processes^[16]^.

One process that has been implicated in a range of psychiatric disorders is the discounting of reward value over time (temporal discounting): both steep and shallow discounting is associated with different psychiatric conditions^[9]^. In temporal discounting tasks, participants make repeated choices between a fixed immediate reward and larger but temporally delayed rewards^[17]^. Based on binary choices and/or response time (RT) distributions, the degree to which participants discount the value of future rewards based on the temporal delay provides a measure of individual impulsivity. Increased temporal discounting is thought to be a trans-diagnostic marker with relevance for a range of psychiatric disorders^[9]^, with addictions and related disorders being prominent examples^[18,19]^.

There is preliminary evidence that temporal discounting might be more pronounced when addiction related cues are present. Participants who suffer from gambling disorder for instance tend to exhibit steeper discounting ^[12,20]^ and increased risk-taking^[21]^ in the presence of gambling-related stimuli or environments. These findings resonate with theories of drug addiction such as incentive sensitization theory^[22]^ which emphasize a prominent role for addiction-related cues in the maintenance of drug addiction (see below). Identifying the mechanisms underlying such behavioral patterns and how they are modulated by addiction-related cues is essential to the planning and execution of successful interventions that aim to reverse these changes in decision-making^[23,24]^.

Accordingly, the concept of cue-reactivity plays a prominent role in research on substance use disorders^[25]^, but has more recently also been investigated in behavioral addictions such as gambling disorder^[26]^. Cue-reactivity refers to conditioned responses to addiction-related cues in the environment and is thought to play a major role in the maintenance of addiction. Cue-reactivity can manifest in behavioral measures, as described above for temporal discounting and risk-taking, but also in subjective reports and/or in physiological measures^[25]^. Incentive-Sensitization Theory^[22,27]^ states that neural circuits mediating the incentive motivation to obtain a reward become over-sensitized to addiction-related cues, giving rise to craving. These motivational changes are thought to be mediated by dopaminergic pathways of the mesocorticolimbic system^[28–30]^. In line with this, craving following cue exposure correlates with a modulation of striatal value signals during temporal discounting^[12]^, and exposure to drug-related cues increases dopamine release in striatal circuits in humans^[30]^. While studying these mechanisms in substance use disorders is certainly of value, it is also problematic because substances might have direct effects on the underlying neural substrates. Behavioral addictions, such as gambling disorder, however, might offer a somewhat less perturbed view on the underlying mechanisms.

Studies probing cue-reactivity in participants suffering from gambling disorder have typically either used picture stimuli^[12,13,21,31–38]^ or real-life gambling environments (i.e. gambling facilities)^[20]^. Both methods come with advantages and disadvantages. While presenting pictures in a controlled lab environment enables researchers to minimize the influence of noise factors and simplifies the assessment physiological variables, it lacks the ecological validity of real-life environments. Conversely, a field study in a real gambling outlet arguably has high ecological validity but lacks the control of confounding factors and makes it difficult to obtain physiological measures.

By equipping participants with head-mounted VR-glasses and sufficient space to navigate within the VR-environment, a strong sense of immersion can be created, which in turn generates more realistic stimulation. In this way VR also offers a potential solution for the problem of ecologically valid addiction-related stimuli for studies in the field of cue-reactivity^[7,8]^. For example, Bouchard et al.^[2]^ developed a VR-design that is built to provide ecologically valid stimuli for participants suffering from gambling disorder by placing them in a virtual casino. The design can be used in treatment in order to test reactions and learned cognitive strategies in a secure environment. The present study builds upon this idea to create a design that allows assessment of behavioral, subjective and physiological cue-reactivity in VR-environments. Participants are immersed in two rich and navigable VR environments that either represent a (neutral) café environment or a gambling-related casino environment. Within these environments, behavioral cue-reactivity can be measured via behavioral tasks implemented in VR. Given that immersion in the virtual environment takes place in a controlled lab setting, the measurement of physiological variables like electrodermal activity^[39]^ and heart rate, as indicators of physiological cue-reactivity^[25,26]^, is also easily accommodated.

Studies using computational modeling to asses latent processes underlying learning and decision-making increasingly include not only binary decisions, but also response times (RTs) associated with these decisions, e.g. via sequential sampling models such as the drift diffusion model (DDM)^[40]^. This approach has several potential advantages. First, leveraging the information contained in the full RT distributions can improve the stability of parameter estimates^[41,42]^. Second, by conceiving decision making as a dynamic diffusion process, a more detailed picture of the underlying latent processes emerges^[43–47]^. Recent studies, for instance, applied these techniques to temporal discounting, where they revealed novel insights into effects of pharmacological manipulation of the dopamine system on choice dynamics^[46]^. Likewise, we applied these techniques to examine the processes underlying reinforcement learning impairments in gambling disorder^[48]^ and decision-making alterations following medial orbitofrontal cortex lesions^[45]^. Importantly, most standard lab-based testing settings use keyboards, button boxes and computer screens to record responses and display stimuli during behavioral tasks. In contrast, in the present study we used VR-controllers in a 3D virtual space. This represents a fundamentally different response mode, because in VR, participants have to physically move the controller to the location of the chosen option and then execute a button press to indicate their choice, adding additional motor complexity. In particular in the context of RT-based modeling, a crucial question is therefore whether responses obtained via VR-controllers allow for a comprehensive RT-based computational modeling, as previously done using standard approaches. Therefore, we also explored the applicability of drift diffusion modeling in the context of behavioral data obtained in VR.

Besides validating our VR-design with a healthy cohort of participants, the study at hand investigated the stability of parameters derived from temporal discounting tasks, in particular the discount rate *log(k)*. Recently, the reliability of behavioral tasks as trait indicators of impulsivity and cognitive control has been called into question^[49,50]^, in particular when compared to questionnaire-based measures of self-control^[49]^. It has been argued that the inherent property that makes behavioral tasks attractive for group-based comparisons renders them less reliable as trait markers^[51]^. Specifically, Hedge et al.^[51]^ argue that tasks having a low between participant variability produce robust group effects in experimental studies and are therefore employed frequently. However, some of these tasks suffer from reduced test-retest-reliability for individual participants due to their low between-participant variability. Notably, Enkavi et al.^[49]^ reported a reliability of .65 for the discount rate *k*, the highest of all behavioral tasks examined in that study, and comparable to the reliability estimates of the questionnaire-based measures. This is in line with previous studies on the reliability of *k*, which provided estimates ranging from .7 to .77^[52,53]^. Importantly, as outlined above, both the actual response mode and the contextual setting of VR-based experiments differ substantially from standard lab-based testing situations employed in previous reliability studies of temporal discounting^[49,52–55]^. Therefore, it is an open question whether temporal discounting measures obtained in VR exhibit a reliability comparable to the standard lab-based tests that are typically used in psychology.

Taken together, by examining healthy non-gambling participants on different days and under different conditions (neutral vs. gambling-related VR environment, standard lab-based testing situation), we addressed the issue of reliability of temporal discounting in virtual vs. standard lab environments. We furthermore explored the feasibility of applying the drift diffusion model in the context of RTs obtained via VR-compatible controllers. Finally, we also examined physiological reactivity during exploration of the different virtual environments. The specific virtual environments employed here are ultimately aimed to examine these processes in gambling disorder (e.g. the setup includes a gambling-related and a neutral cafe environment). However, the present study has more general implications for the application of behavioral and psychophysiological testing in virtual environments by examining the reliability of model-based analyses of decision-making in lab-based testing vs. testing in different VR environments in a group of young non-gambling controls.

We hypothesized that the data produced on different days and under different conditions would yield only little evidence in favor of systematic shifts in temporal discounting behavior within a group of healthy non-gambling participants, suggesting only insubstantial effects caused by the different environments in our VR-design. Furthermore, we hypothesized that temporal discounting would show a strong reliability, adding further strength to the case that temporal discounting is stable over time and can be applied in VR. Finally, we hypothesized that we could capture latent decision variables in a VR context with the DDM.

## Methods

### Participants

Thirty-four healthy participants (25 female) aged between 18 and 44 (mean = 26.41, std = 6.44) were invited to the lab on three different occasions. Participants were recruited via flyers at the University of Cologne and via postings in local internet forums. No participant indicated a history of traumatic brain injury, psychiatric or neurological disorders or severe motion sickness. Participants were additionally screened for gambling behavior using the questionnaire Kurzfragebogen zum Glückspielverhalten (KFG)^[56]^. The KFG fulfills the psychometric properties of a reliable and valid screening instrument. No participant showed a high level (>15 points on the KFG) of gambling affinity (mean = 1.56, std = 2.61, range: 0 to 13).

Participants provided informed written consent prior to their participation, and the study procedure was approved by the Ethics Board of the Germany Psychological Society. The procedure was in accordance with the 1964 Helsinki declaration and its later amendments or comparable ethical standards.

### VR-Setup

The VR-environments were presented using a wireless HTC VIVE head-mounted display (HMD). The setup provided a 110° field of view, a 90 Hz refresh rate and a resolution of 1440 x 1600 Pixel per eye. Participants had an area of about 6m^2^ open space to navigate the virtual environment. For the execution of the behavioral tasks and additional movement control participants held one VR-controller in their dominant hand. The VR-software was run on a PC with the following specifications: CPU: Intel Core i7-3600, Memory: 32.0 GB RAM, Windows 10, GPU: NVIDIA GeForce GTX 1080 (Ti). The VR-environments themselves were designed in Unity. Auditory stimuli were presented using on-ear headphones.

### VR-Environments

The two VR-environments both consisted of a starting area and an experimental area. The starting area was the same for both VR-environments. It consisted of a small rural shopping street and a small park. Participants heard low street noises. The area was designed for familiarization with the VR-setup and the initial exploration phase. The experimental area of the environments differed for the two environments. For the VR_neutral_ environment it contained a small café with a buffet (Figure 1 a, b and c). Participants could hear low conversations and music. The gambling-related environment (VR_gambling_) contained a small casino with slot machines and a sports betting area (Figure 1 d, e and f). The audio backdrop was the sound of slot machines and sports. The floorplan of both of these experimental areas was identical but mirrored for the café (Figure 1 a and d). Both experimental areas additionally included eight animated human avatars. These avatars performed steady and non-repetitive behaviors like gambling and ordering food for the gambling-related and neutral environments, respectively. Both experimental areas (café and casino) had entrances located at the same position within the starting area of the VR-environments, which were marked by corresponding signs.

**Figure 1.**
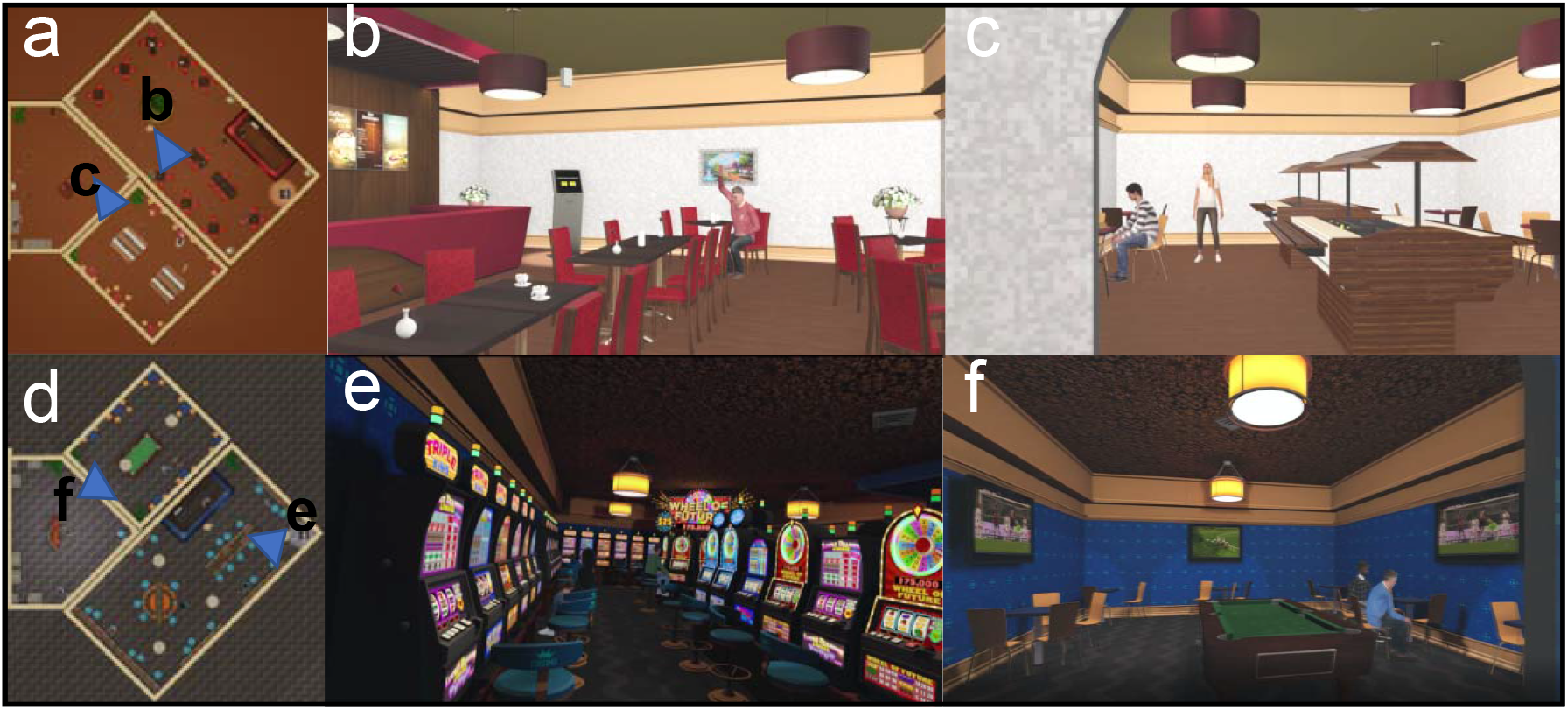
Experimental areas of the VR-environments a) Floorplan of the café within the VR-neutral environment b) View of the main room of the café c) View of the buffet area of the café d) Floorplan of the casino within the VR-gambling environment e) View of the main room of the casino f) View of the sports bar within the casino

### Experimental procedure

Participants were invited to the VR lab for three different sessions on three different days. The time between the sessions was between one day and nineteen days (mean = 3.85, std = 3.36). During the three sessions participants either explored one of two different VR environments (VR-sessions) followed by the completion of two behavioral tasks, or simply performed the same two behavioral tasks in a standard lab-testing context (Lab-session). If the session was a VR-session, electrodermal activity (EDA)^[39]^ was measured during a non-VR baseline period and the exploration of the VR-environments. The order of the sessions was pseudorandomized. At the first session, not depending on if VR was applied or not, participants arrived at the lab and the behavioral tasks were explained in detail. If the session was a Lab-session, participants proceeded with the two behavioral tasks. If the session was the first of the VR-sessions, participants were subsequently familiarized with the VR-equipment and handling. Participants were seated and a five-minute EDA baseline was measured (baseline phase). For both VR-sessions participants were then helped to apply the VR-equipment and entered the VR-environments. Within the VR-environments participants first explored the starting area for 5 minutes (first exploration phase). After these five minutes participants were asked to enter the experimental area of the environment (either the café or the casino) (Figure 1). Participants were instructed to explore the interior experimental area for five minutes (second exploration phase). Each of the three phases was later binned into five one-minute intervals and labeled as B (1 to 5) for the baseline phase, F (1 to 5) for the first exploration phase and S (1 to 5) for the second exploration phase. During the exploration the experimenter closely monitored the participants and alerted them if they were about to leave the designated physical VR-space. After the second exploration phase participants were asked to proceed to a terminal within the VR-environment on which the behavioral tasks were presented.

### Physiological measurements

EDA was measured using a BioNomadix-PPGED wireless remote sensor together with a Biopac MP160 data acquisition system (Biopac Systems, Santa Barbara, CA, USA). A GSR100C amplifier module with a gain of 5V, low pass filter of 10 Hz and a high pass filter DC were included in the recording system. The system was connected to the acquisition computer running the AcqKnowledge software. Triggers for the events within the VR-environments were send to the acquisition PC via digital channels from the VR-PC. Disposable Ag/AgCl electrodes were attached to the thenar and hypothenar eminences of the non-dominant palm. Isotonic paste (Biopac Gel 101) was used to ensure optimal signal transmission. The signal was measured in micro-Siemens units (mS).

### Behavioral Tasks

Participants performed the same two behavioral tasks with slightly varied rewards and choices in each of the three sessions: a temporal discounting task^[17]^ and a 2-step sequential decision-making task^[57,58]^. Results from the 2-step task will be reported separately. In the temporal discounting task participants had to repeatedly choose between an immediately available (smaller-but-sooner, SS) monetary reward of 20 Euros and larger-but-later (LL) temporally delayed monetary rewards. The LL options were multiples of the SS option (range 1.025 to 3.85) combined with different temporal delays (range 1 to122 days). We constructed three sets of six delays and 16 LL options. Each set had the same mean delay and the same mean LL option. Combining each delay with every LL option within each set resulted in three sets of 96 trials. The order of presentation of the trial sets was counter balanced across participants and sessions. All temporal discounting decisions were hypothetical^[59,60]^. In the VR-version of the task two yellow squares were presented to the participants (Figure 2). One depicted the smaller offer of 20 Euros now, while the other depicted the delayed larger offer. For the lab-based testing session were presented in the same way except that the color scheme was white writing on a black background. Offers were randomly assigned to the left/right side of the display and presented until a decision was made. The next trial started .5 to 1 seconds after the decision. Participants indicated their choice either by aiming the VR-controller at the preferred option and pulling the trigger (VR-sessions) or by pressing the corresponding arrow key on the keyboard (Lab-session).

**Figure 2.**
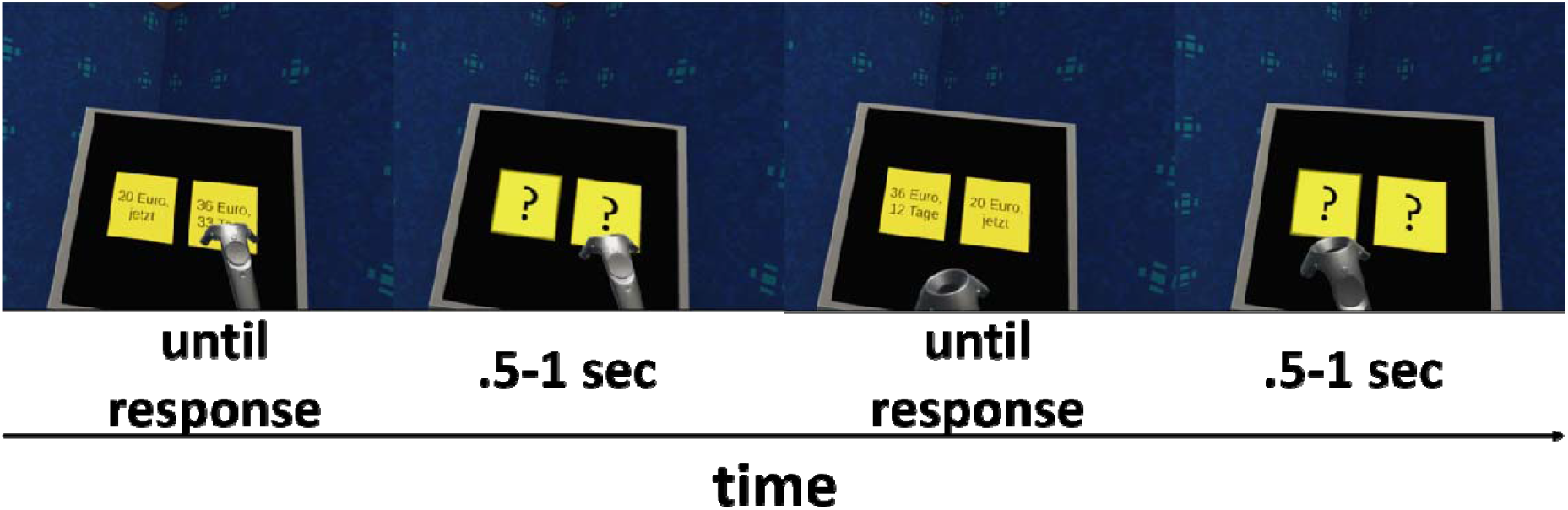
Presentation of the temporal discounting task in VR. Participants had to repeatedly decide between a small but immediate reward (SS) and larger but temporally delayed rewards (LL). Amounts and delays were presented in yellow squares. During the inter-trial intervals (.5-1 sec.) these squares contained only question marks. Participants indicated their choice by pointing the VR-controller at one of the yellow squares and pulling the trigger.

### Model-free discounting data analysis

The behavioral data from the temporal discounting task was analyzed using several complementary approaches. First, we used a model-free approach that involved no a priori hypotheses about the mathematical shape of the discounting function. For each delay, we estimated the LL reward magnitudes at which the subjective value of the LL reward was equal to the SS (indifference point). This was done by fitting logistic functions to the choices of the participants, separately for each delay. Subsequently, these indifference points were plotted against the corresponding delays, and the area under the resulting curve (AUC) was calculated using standard procedures^[61]^. AUC values were derived for each participant and testing session, and further analyzed with the intra-class correlation (ICC) and the Friedman Test, a non-parametric equivalent of the repeated measures ANOVA model.

### Computational modeling

Previous research on the effects of the delay of a reward on its valuation proposed a hyperbolic nature of devaluation^[62,63]^. Therefore, the rate of discounting for each participant was also determined employing a cognitive modeling approach using hierarchical Bayesian modeling^[16]^. A hierarchical model was fit to the data of all participants, separately for each session (see below). We applied a hyperbolic discounting model (equation 1):

Here, SV(LL) denotes the subjective (discounted) value of the LL. A and D represent the amount and the delay of the LL, respectively. The parameter k governs the steepness of the value decay over time, with higher values of k indicating steeper discounting of value over time. As the distribution of the discount rate k is highly skewed, we estimated the parameter in log-space (log[k]), which avoids numerical instability in estimates close to 0.

The hyperbolic model was then combined with two different choice rules, a softmax action selection rule^[64]^ and the drift diffusion model^[44]^. For softmax action selection, the probability of choosing the LL option on trial *t* is given by equation (2).

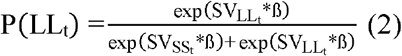

Here, the ß-parameter determines the stochasticity of choices with respect to a given valuation model. A ß of 0 would indicate that choices are random, whereas higher ß values indicate a higher dependency of choices on option values. The resulting best fitting parameter estimates were used to test the ICC and systematic session effects via comparison of the posterior probabilities of group parameters.

Next, we incorporated response times (RTs) into the model by replacing the softmax choice rule with the drift diffusion model (DDM)^[43–46]^. The DDM models choices between two options as a noisy evidence accumulation that terminates as soon as the accumulated evidence exceeds one of two boundaries. In this analysis the upper boundary was set to represent LL choices, and the lower boundary SS choices. RTs for choices of the immediate reward were multiplied by −1 prior to model estimation. To prevent outliers in the RT data from negatively impacting model fit, the 2,5% slowest and fastest trials of each participant were excluded from the analysis^[44,45]^. In the DDM the RT on trial *t* is distributed according to Wiener first passage time (wfpt) (equation 3).

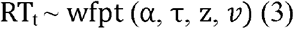

Here α represents the boundary separation modeling the tradeoff between speed and accuracy. τ represents the non-decision time, reflecting perception and response preparation times. The starting value of the diffusion process is given by z, which therefore models a potential bias towards one of the boundaries. Finally, rate of evidence accumulation is given by the drift-rate *v*.

We first fit a null model (DDM_0_), where the value difference between the two options was not included, such that DDM parameters were constant across trials^[45,46]^. We then used two different temporal discounting DDM_S_, in which the value difference between options modulated trial-wise drift rates. This was done using either a linear (DDM_L_) or a non-linear sigmoid (DDM_S_) linking function^[47]^. In the DDM_L,_ the drift-rate v in each trial is linearly dependent on the trial-wise scaled value difference between the LL and the SS options (equation 4) ^[44]^. The parameter v_coeff_ maps the value differences onto v and scales them to the DDM:

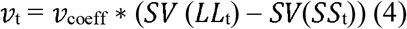

One drawback of a linear representation of the relationship between the drift-rate v and trial-wise value differences is that *v* might increase infinitely with high value differences, which can lead the model to under-predict RTs for high value differences^[45]^. In line with previous work ^[45,46]^ we thus included a third version of the DDM, that assumes a non-linear sigmoidal mapping from trial-wise value differences to drift rates (equations 5 and 6)^[43]^:

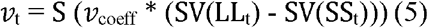

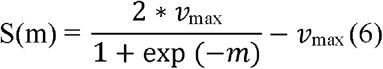

Here, the linear mapping function from the DDM_L_ is additionally passed through a sigmoid function S with the asymptote v_max_, causing the relationship between v and the scaled trail-wise value difference *m* to asymptote at v_max_.

We have previously reported detailed parameter recovery analyses for the DDM_S_ in the context of value-based decision-making tasks such as temporal discounting^[45]^, which revealed that both subject-level and group-level parameters recovered well.

### Hierarchical Bayesian Models

All models were fit to the data of all participants in a hierarchical Bayesian estimation scheme, separately for each session, resulting in independent estimates for each participant per session. Participant-level parameters were assumed to be drawn from group-level Gaussian distributions, the means and precisions of which were again estimated from the data. Posterior distributions were estimated via Markov Chain Monte Carlo in the R programming language^[65]^ using the JAGS software package^[66]^. For the DDM’s the Wiener module for JAGS was used^[67]^. For the group-level means, uniform priors over numerically plausible parameter ranges were chosen (Table 1). Priors for the precision of the group-level distribution were Gamma distributed (0.001, 0.001). The convergence of chains was determined by the R-hat statistic^[68]^. Values between 1 and 1.01 were considered acceptable. Comparisons of relative model fit were performed using the Deviance Information Criterion (DIC), where lower values reflect a superior model fit^[69]^.

**Table 1.** Ranges for the uniform priors of group-level parameter means. Ranges were chosen to cover numerically plausible values. Parameters included in multiple models are only listed once.

| Parameter | Prior for group mean |
| --- | --- |
| **log(k)** | *Uniform*(-20, 3) |
| **softmax $\beta$** | *Uniform*(0, 10) |
| **$\nu$** | *Uniform*(-100, 100) |
| **$\tau$** | *Uniform*(.1, 6) |
| **$\alpha$** | *Uniform*(.01, 5) |
| **$z$** | *Uniform*(.1, .9) |
| **$\nu_{\text{coeff}}$** | *Uniform*(-100, 100) |
| **$\nu_{\text{max}}$** | *Uniform*(0, 100) |

### Systematic session effects on model parameters

Potential systematic session effects on group level posterior distributions of parameters of interest were analyzed by overlaying the posterior distributions of each group level parameter for the different sessions. Here we report the mean of the posteriors of the estimated group level parameters and the difference distributions between them, the 95% highest density intervals (HDI) for both of these as well as directional Bayes Factors (dBF) which quantify the degree of evidence for reductions vs. increases in a parameter. Because the priors for the group effects are symmetric, this dBF can simply be defined as the ratio of the posterior mass of the difference distributions above zero to the posterior mass below zero^[70]^. Here directional Bayes Factors above 3 are interpreted as moderate evidence in favor of a positive effect, while Bayes Factors above 12 are interpreted as strong evidence for a positive effect^[71]^. Specifically, a dBF of 3 would imply that a positive directional effect is three times more likely than a negative directional effect. Bayes Factors below 0.33 are likewise interpreted as moderate evidence in favor of the alternative model with reverse directionality. A dBF above 100 is considered extreme evidence^[71]^. The cutoffs used here are liberal in this context, because they are usually used if the test is against a H_0_ implying an effect of 0. In addition, we report the effect size (Cohen’s *d*) based on the mean posterior distributions of the session means, the pooled standard deviations across sessions and the correlation between sessions.

### ICC analysis

The test-retest reliability of the best fitting parameter values between the three sessions was analyzed using the intra-class correlation coefficient (ICC). The ICC-analysis was done in the R programming language^[65]^ and was based on a mean-rating of three raters, absolute agreement and a two-way mixed model. ICC values below .5 are an indication of poor test-retest reliability, whereas values in the range between .5 and .75 indicate a moderate test-retest reliability^[72]^. Higher values between .75 and .9 indicate a good reliability, while values above .9 suggest an excellent test-retest reliability.

### Analysis of physiological data

A frequently used index of sympathetic activity is electrodermal activity, i.e. changes in skin conductance (SC)^[73]^. Here the physiological reactivity to the VR-environments is measured as the slowly-varying skin conductance level (SCL)^[39]^. Thus, the SCL was extracted from the EDA signal using continuous decomposition analysis (CDA) via the Ledalab toolbox^[74]^ for Matlab (MathWorks). For the deconvolution, default settings were used. The resulting signal was then transformed into percentage change from the mean signal of the five minutes baseline phase at the beginning of the experiment. Subsequently, five one-minute bins were constructed for each phase of the VR-session (baseline phase, the first exploration phase and the second exploration phase). An alternative way of classifying tonic sympathetic arousal can be the number of spontaneous phasic responses (SCR) in the EDA signal^[74]^. Again, the signal was divided in one-minute bins and the number of spontaneous SCRs during each bin was calculated from the phasic component of the deconvoluted EDA signal using the Ledalab toolbox. The resulting values were similarly transformed into percentage change from the mean number of SCRs during the five baseline bins. To test whether entering the VR-environments had a general effect on sympathetic arousal, we compared the values for the last time point of the base line phase (B5) with the first time point of the first exploration phase (F1) for both sessions using a non-parametric Wilcoxon Signed-Rank Test. To test whether there was a differential effect of entering the different experimental areas of the VR-environments on sympathetic arousal, for both measures the differences between the last time point of the first exploration phase (F5) and the first time point of the second exploration phase (S1) were compared across VR-sessions using a non-parametric Wilcoxon Signed-Tanks Test^[75]^. Effect sizes are given as *r*^[76]^, computed as the statistic Z divided by the square-root of N. Effect sizes between 0 and .3 are considered small and effect sizes between .3 and .5 are considered medium and *r* values > .5 are considered large effects.

### Data and code availability

Raw behavioral and physiological data as well as JAGS model code is available on the Open Science Framework (https://osf.io/xkp7c/files/).

## Results

### Temporal discounting AUC

The analysis of the AUC values revealed no significant session effect across participants (Friedman Test: Chi-Squared = 1.235 df =2 p =.539). Furthermore, the ICC value was .93 (95% confidence interval (CI): .89 - .96) (p<.001) indicating an excellent test-retest reliability of temporal discounting AUC values over the three sessions (Table 2). Pairwise correlations between all sessions can be found in the supplementary materials (Supplementary Figure S1).

**Table 2.**
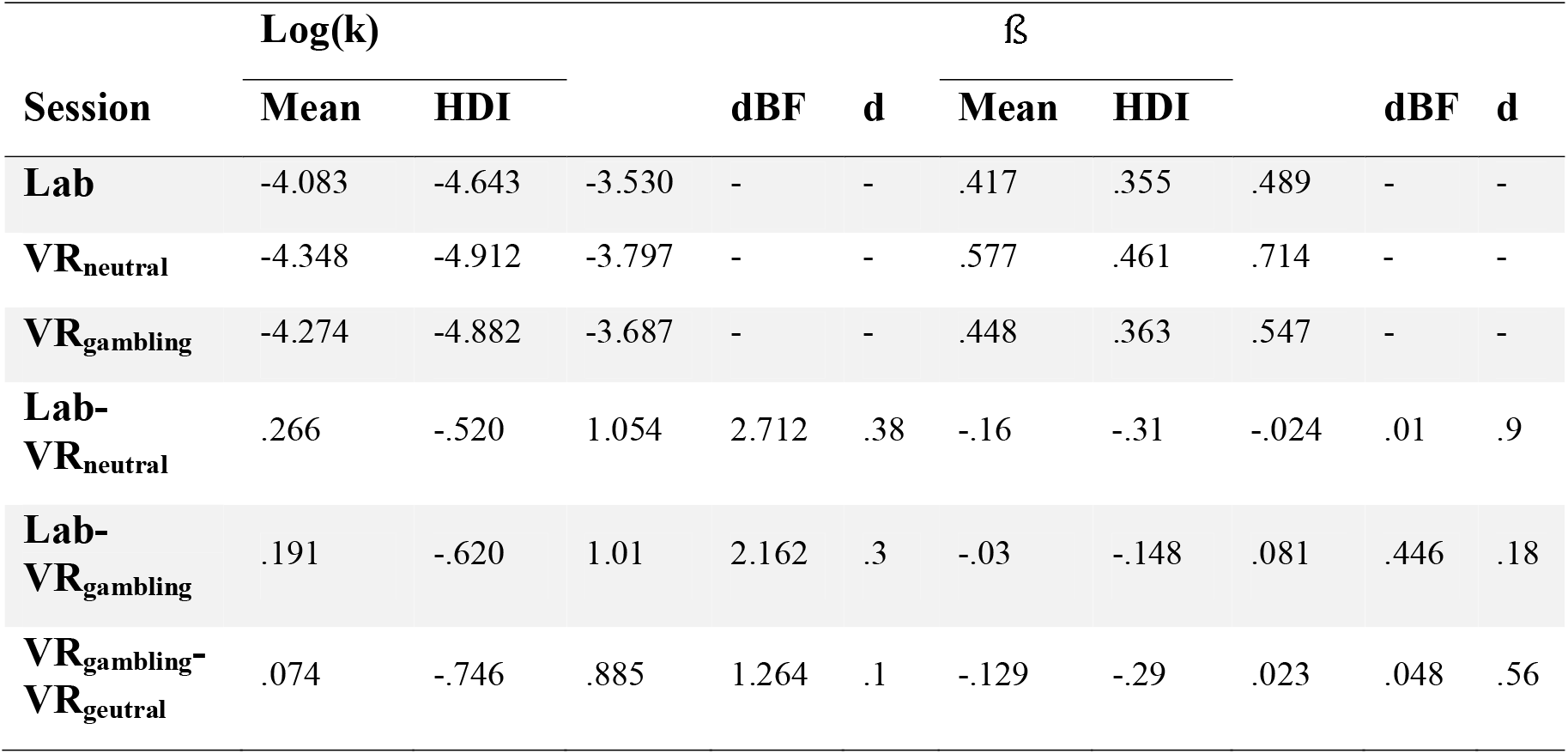
95% HDIs for the two parameters of the hyperbolic discounting model. HDIs are described by the min. value first and the max value second. Directional Bayes Factors (dBF) are calculated as BF = i/(1-i), with i being the probability mass of the difference distributions above zero. Effect sizes are given as Cohen’s d.

### Softmax choice rule

For the hyperbolic model with softmax choice rule, the group level posteriors showed little evidence for systematic effects of the different sessions on log(k) (all BFs < 3 or >.33) (Figure 3a and c and Table 2). In contrast, the softmax f3 parameter was higher (reflecting higher consistency) in the VR_neutral_ session compared to the other sessions (vs. Lab: dBF = .01 and vs. VR_gambiling_: dBF = .048) (Figure 3b and d, Table 2). This indicates that a higher f3 in the VR_neutral_ session was approximately 100 (Lab) or 20 (VR_gambling_) times more likely than a lower f3. There was little evidence for a systematic effect between the Lab and VR_gambling_ sessions (dBF = .446).

**Figure 3.**
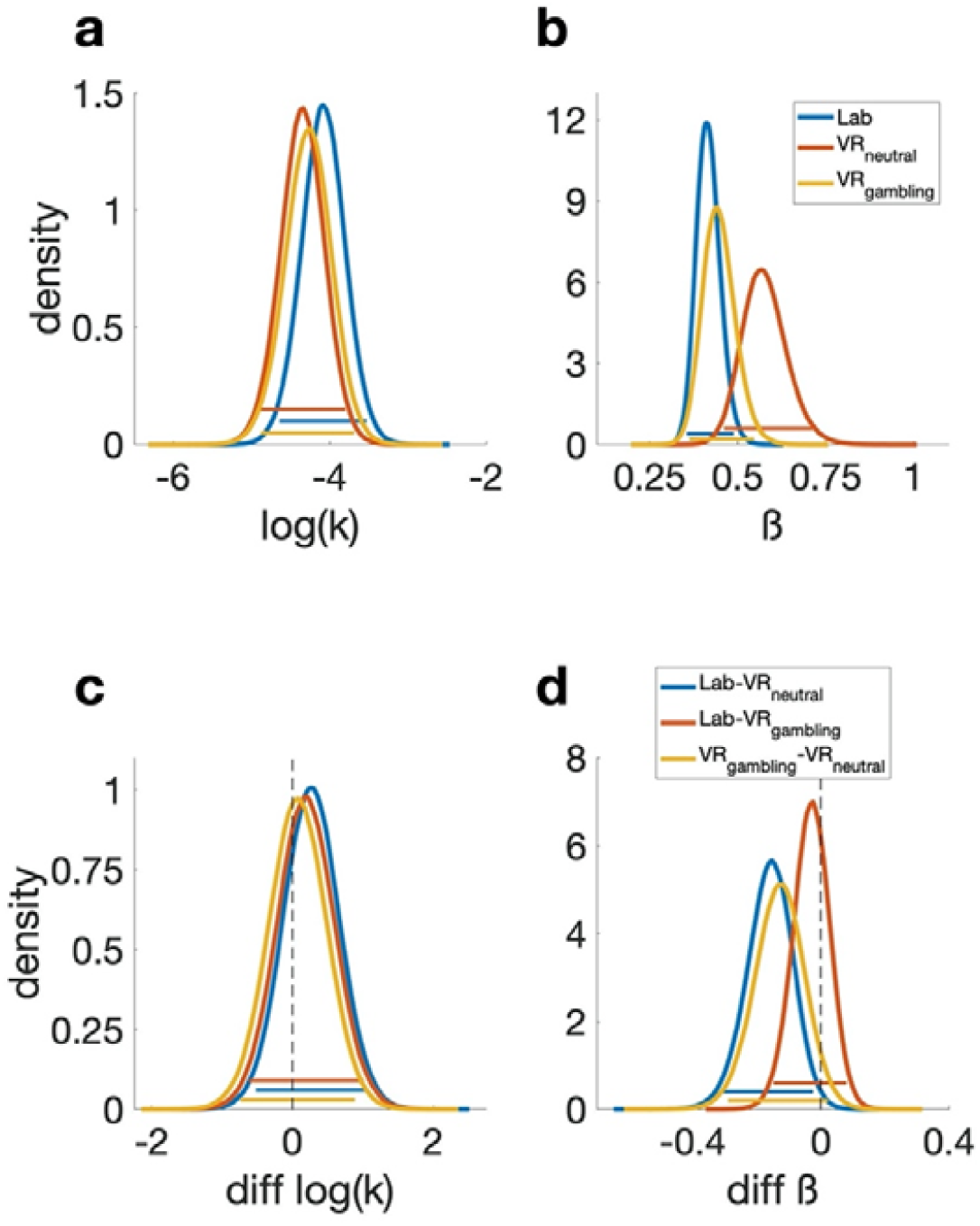
Posterior distributions of the parameters of the hyperbolic discounting model. Colored bars represent the corresponding 95% HDIs. a) Posterior distribution of the log(k) parameter (reflecting the degree of temporal discounting) for all three sessions. b) Posterior distribution of the *β* or inverse temperature parameter (reflecting decision noise). c) Pairwise difference distributions between the posteriors of the log(k) parameters of all three sessions. d) Pairwise difference distributions between the posteriors of the *β* parameters of all three sessions.

The ICC value for the log(k) parameter indicated an excellent test-retest reliability of .91 (CI: .86 - .96) (p<.001) (Table 3). For the *β*-parameter of the softmax choice rule the ICC value was .34 (CI: .17 - .53) (p<.001) indicating a poor test-retest reliability (Table 3). The pairwise correlations of estimated parameter values between all sessions can be found in the Supplement (Supplementary Figure S2 and S3). Pairwise correlations between all sessions for both parameters can be found in the supplementary materials (Supplementary Figure S2 and S3).

**Table 3.** Summary of the results of the ICC analysis for the AUC values as well as the two parameters of the hyperbolic discounting model with a softmax choice rule. Lower and upper bound describe the 95% confidence interval.

| Parameter | ICC | p | Lower bound | Upper Bound |
| --- | --- | --- | --- | --- |
| AUC | .93 | <.001 | .89 | .96 |
| $\log(k)$ | .91 | <.001 | .86 | .95 |
| $\beta$ | .34 | <.001 | .17 | .53 |

### Drift diffusion model choice rule

Model comparison revealed that the DDM_S_ had the lowest DIC in all conditions (Table 4) replicating previous work ^[45,46,48]^. Consequently, further analyses of session effects and reliability focused on this model. For the log(k) parameter, the 95% HDIs showed a high overlap between all sessions indicating no systematic session effects, however the BFs showed moderate evidence for a reduced log(k) in the VR_neutral_-session (Figure 4 a and d, Table 5). A lower value in the VR_neutral_-session was about seven (Lab-session dBF = 6.756) or four times (VR_gambling_ dBF = 3.86) more likely than a lower value. Similarly, the posterior distributions of *v*_max_, *v*_coeff_ and α were highly overlapping, whereas some of the dBFs gave moderate evidence for systematic directional effects within these parameters (Figure 4 b, c, e and f, Figure 5 b and e, Table 5). v_coeff_, mapping trial-wise value difference onto the drift rate, was lowest in the Lab-session and highest in VR_neutral_ (Lab-VR_neutral_ dBF = .074, Lab-VR_gambling_ = .2, VR_gambling_-VR_neutral_ = .228). Thus, an increase in *v*_coeff_ in VR_neutral_ compared to the Lab-session was approximately thirteen times more likely than a decrease. Likewise, it was approximately five times more likely that there was an increase in the VR_neutral_ compared to the VR_gambling_-session. For v_max_, the upper boundary for the value difference’s influence on the drift rate, the dBFs indicated that a positive shift from VR_gambling_ to VR_neutral_ was five times more likely than a negative shift (Dbf = .203) but there was only very little indication of a systematic difference between both of them and the Lab-session. Finally, a reduction of the boundary separation parameter a was five times more likely than an increase when comparing the VR_neutral_ to the Lab-session (Dbf = .255). There was little evidence for any other systematic differences. The bias parameter *z* displayed high overlap in HDIs and little evidence for any systematic effects between sessions (all dBFs >.33 or <3) (Figure 5 c and f, Table 5). For the non-decision time parameter τ there was extreme evidence for an increase in the VR-sessions compared to the Lab-session (both dBFs >100), reflecting prolonged motor and/or perceptual components of the RT that was more than 100 times more likely than a shortening of these components (Figure 5 a and d, Table 5).

**Table 4.** Summary of the DICs of all DDM models in all sessions. Ranks are based on the lowest DIC in all sessions.

| Model | Lab | VR<sub>neutral</sub> | VR<sub>gambling</sub> | Rank |
| --- | --- | --- | --- | --- |
| DDM<sub>0</sub> | 9275.7 | 9569.8 | 9225.7 | 3 |
| DDM<sub>L</sub> | 7558.9 | 7921.4 | 7663.0 | 2 |
| DDM<sub>S</sub> | 6992.3 | 7327.2 | 7033.1 | 1 |

**Table 5.** Directional Bayes Factors (dBF) and effect sizes (Cohen’s d) for all between session comparisons for all parameters of the DDM_S_. Means and HDIs of the posteriors and difference distributions are summarized in the supplementary materials (Supplementary Table S1). BFs are calculated as BF = i/(1-i), with i being the probability mass of the difference distributions above zero.

| Contrast | log(k) | | $v_{\text{coeff}}$ | | $v_{\text{max}}$ | | $\tau$ | | $\alpha$ | | $z$ | |
| --- | --- | --- | --- | --- | --- | --- | --- | --- | --- | --- | --- | --- |
|  | dBF | d | dBF | d | dBF | d | dBF | d | dBF | d | dBF | d |
| Lab-VR<sub>neutral</sub> | 6.756 | .37 | .074 | .37 | .377 | .2 | >100 | 1.2 | .255 | .224 | .530 | .2 |
| Lab-VR<sub>gambling</sub> | 1.679 | .19 | .200 | .59 | 1.573 | .09 | >100 | 1.5 | .358 | .160 | 1.118 | .04 |
| VR<sub>gambling</sub>-VR<sub>neutral</sub> | 3.860 | .29 | .228 | .27 | .203 | .34 | 3.413 | .17 | .629 | .070 | .458 | .2 |

**Figure 4.**
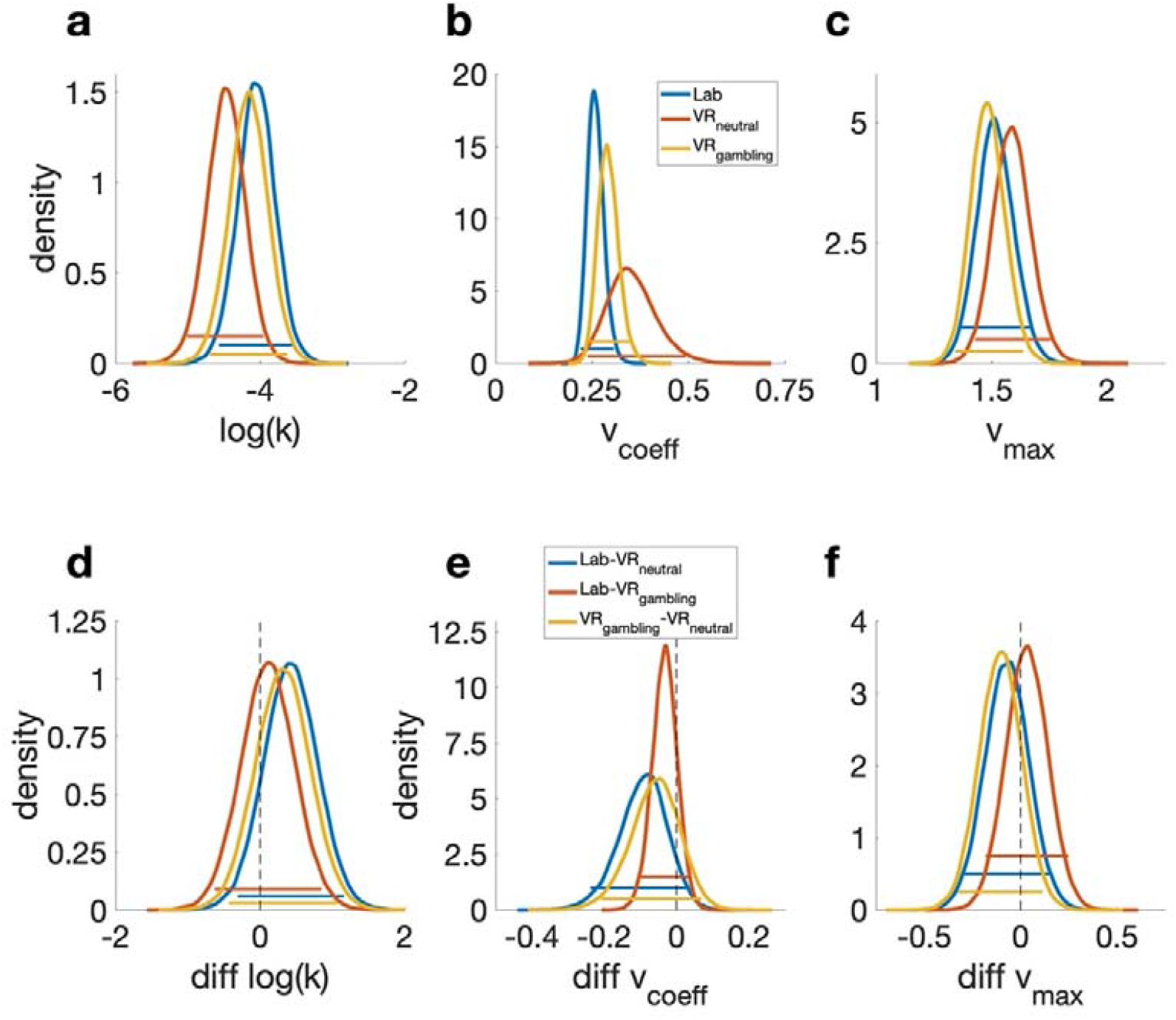
Posterior distributions of the parameters of the DDM_S_ model. Colored bars represent the corresponding 95% HDIs. a) Posterior distributions of the log(k) parameter for all three sessions. b) Posterior distributions of the *v*_coeff_ parameter (mapping the drift rate onto the trial wise value difference). c) Posterior distributions of the *v*_max_ parameter (setting an asymptote for the relation between the trial wise value difference and the drift rate). d) Pairwise difference distributions between the posterior distributions of the log(k) parameters of the three sessions. e) Pairwise difference distributions between the posterior distributions of the *v*_coeff_ parameters of the three sessions. f) Pairwise difference distributions between the posterior distributions of the *v*_max_ parameters of the three sessions.

**Figure 5.**
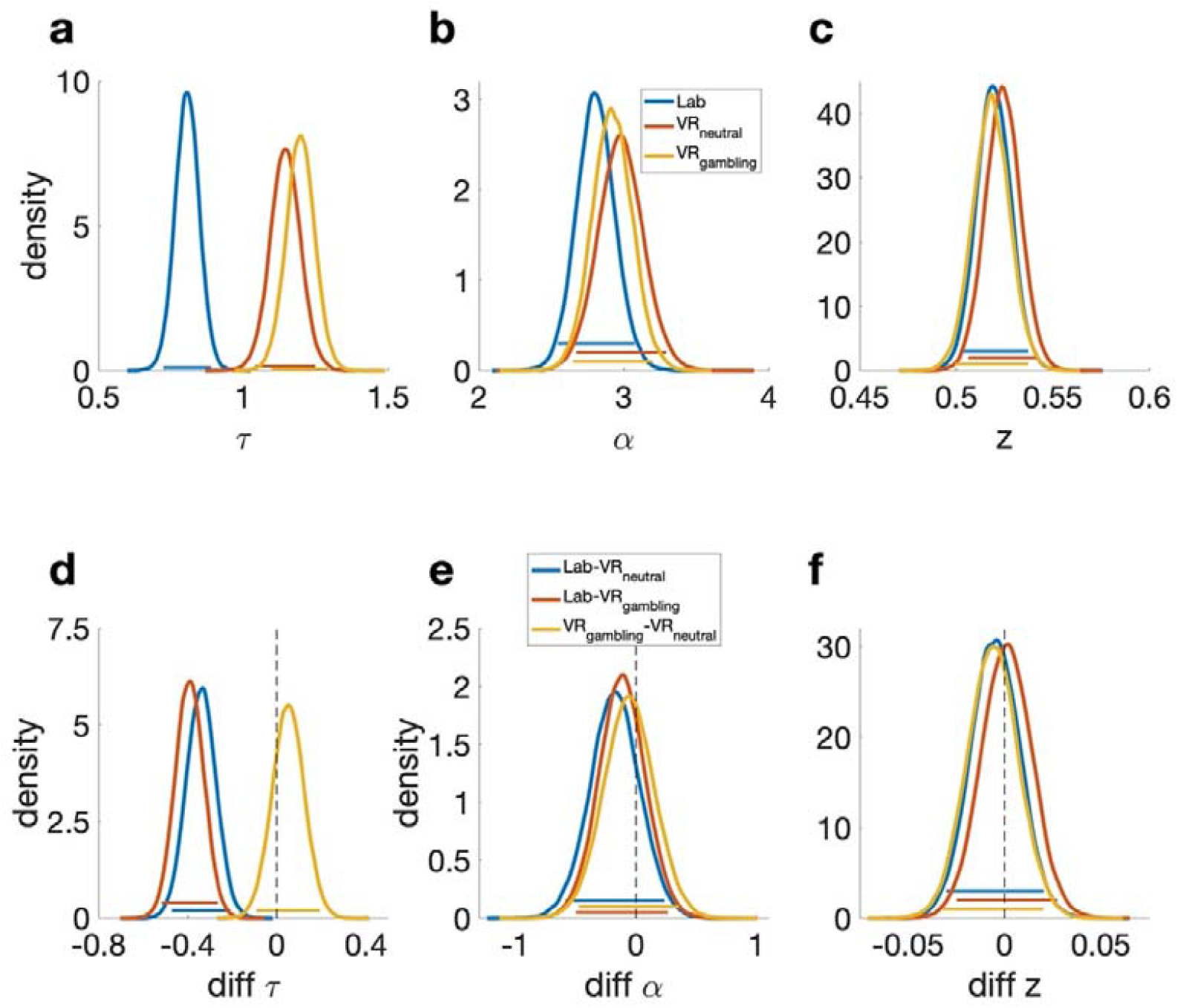
Posterior distributions of the remaining parameters of the DDM_S_ model. Colored bars represent the corresponding 95% HDIs. a) Posterior distributions of the τ parameter (non-decision time) for all three sessions. b) Posterior distributions of the α parameter (separation between decision boundaries). c) Posterior distributions of the z parameter (bias towards one decision option). d) Pairwise difference distributions between the posterior distributions of the τ parameters of the three sessions. e) Pairwise difference distributions between the posterior distributions of the a parameters of the three sessions. f) Pairwise difference distributions between the posterior distributions of the z parameters of the three sessions.

The ICC value for the log(k) parameter was .7 (CI: .56 - .8) indicating a moderate test-retest-reliability (Table 5). For the other DDM_S_ parameters, ICC values were substantially lower (Table 6). Pairwise correlations between all sessions for all parameters can be found in the supplementary materials (Supplementary Figure S4-S9).

**Table 6.** Summary of the results of the ICC analysis of the DDM_S_ parameters.

| Parameter | ICC | p | Lower bound | Upper Bound |
| --- | --- | --- | --- | --- |
| **log(k)** | .7 | <.001 | .56 | .8 |
| **$v_{\text{coeff}}$** | .11 | .14 | -.053 | .3 |
| **$v_{\text{max}}$** | .33 | <.001 | .16 | .52 |
| **$\tau$** | .19 | .033 | .019 | .38 |
| **$\alpha$** | .42 | <.001 | .24 | .59 |
| **$z$** | .4 | <.001 | .22 | .58 |

### Split-half reliability control analyses for DDM parameters

In light of the lower ICC values for the DDM_S_ parameters beyond *log(k)*, we ran additional analyses. Specifically, we hypothesized that these lower ICC values might be attributable to fluctuations of state factors, e.g. mood, fatigue or motivation, between the different sessions. Therefore, we explored within-session reliability of these parameters, separately for each session. Trials where split into odd and even trials and modelled separately using the DDM_S,_ as described above. In general, within-session split-half reliability was substantially greater than test-retest reliability, and mostly in a good to excellent range (range: -.1 for *v*_coeff_ in VR_gambling_ to .94 for τ in VR_neutral_). The lower test-retest reliabilities of some of the DDM_S_ parameters are therefore unlikely to be due to the specifics of the parameter estimation procedure. Rather, these findings are compatible with the view that the parameters underlying the evidence accumulation process might be more sensitive to state-dependent changes in mood, fatigue or motivation. Full results for the split-half reliability analyses can be found in the supplementary materials (Supplementary Tables S3–S5).

### Electrodermal activity (EDA)

The data of 8 of the 34 participants had to be excluded from the EDA analysis, due to technical problems or missing data during one of the testing sessions. Physiological reactivity in the remaining 26 (18 female) participants was analyzed by converting the SCL signal as well as the nSCRs into percent change from the mean level during the base line phase. Both signals were then binned into five one-minute intervals for each of the three phases (baseline, first exploration and second exploration phase). All comparisons were tested with the Wilcoxon Signed Rank Test. Entering the VR-environments (comparing bin B5 to bin F1 for both environments individually) resulted in a significant increase in the SCL values for both VR-environments (VR_neutral_: Z = −3.67, p < .001, r = .72; VR_gambling_: Z = −3.543, p = .002, r = .695) (Figure 6 c and d). The effect was large in both sessions (r > .5). However, for the number of spontaneous SCRs (nSCRs), this effect was only significant in the neutral VR-environment (neutral: Z = −2.623, p = .009, r = .515; gambling: Z = -.013, p = .99, r =.002). There was no significant difference between the two sessions, but the effect was of medium size (Z = −1.7652, p = .078, r =.346) (Figure 6 a and b). To test whether entering the specific experimental areas of the two VR-environments (virtual café vs. virtual casino) had differential effects on physiological responses, the increase in sympathetic arousal from the end of the first exploration phase to the start of the second exploration phase was examined (comparing bin F5 to bin S1, see Figure 6 b and d). The SCL (neutral: Z = - 0.7238, p = -.469, r = .142; gambling: Z = -.089, p = .929, r = .017) as well as the nSCRs (neutral: Z = −1.943, p = .052, r = .381; gambling: Z = .982, p = .326, r = .193) assessed for each session individually showed no significant effect. The effect size was medium (r = .381) for the nSCRs of the VR_neutral_-session and small for all other comparisons (r < .3). Furthermore, the Wilcoxon Signed-Ranks test indicated no significant differences between the two experimental areas on both sympathetic arousal measures (SCL: Z = -.572, p = .381, r = .11; nSCRs.: Z = −1.7652, p = .078, r = .346) (Figure 6 b and d). For the nSCRs however, the effect was of a medium size (r = .346).

**Figure 6.**
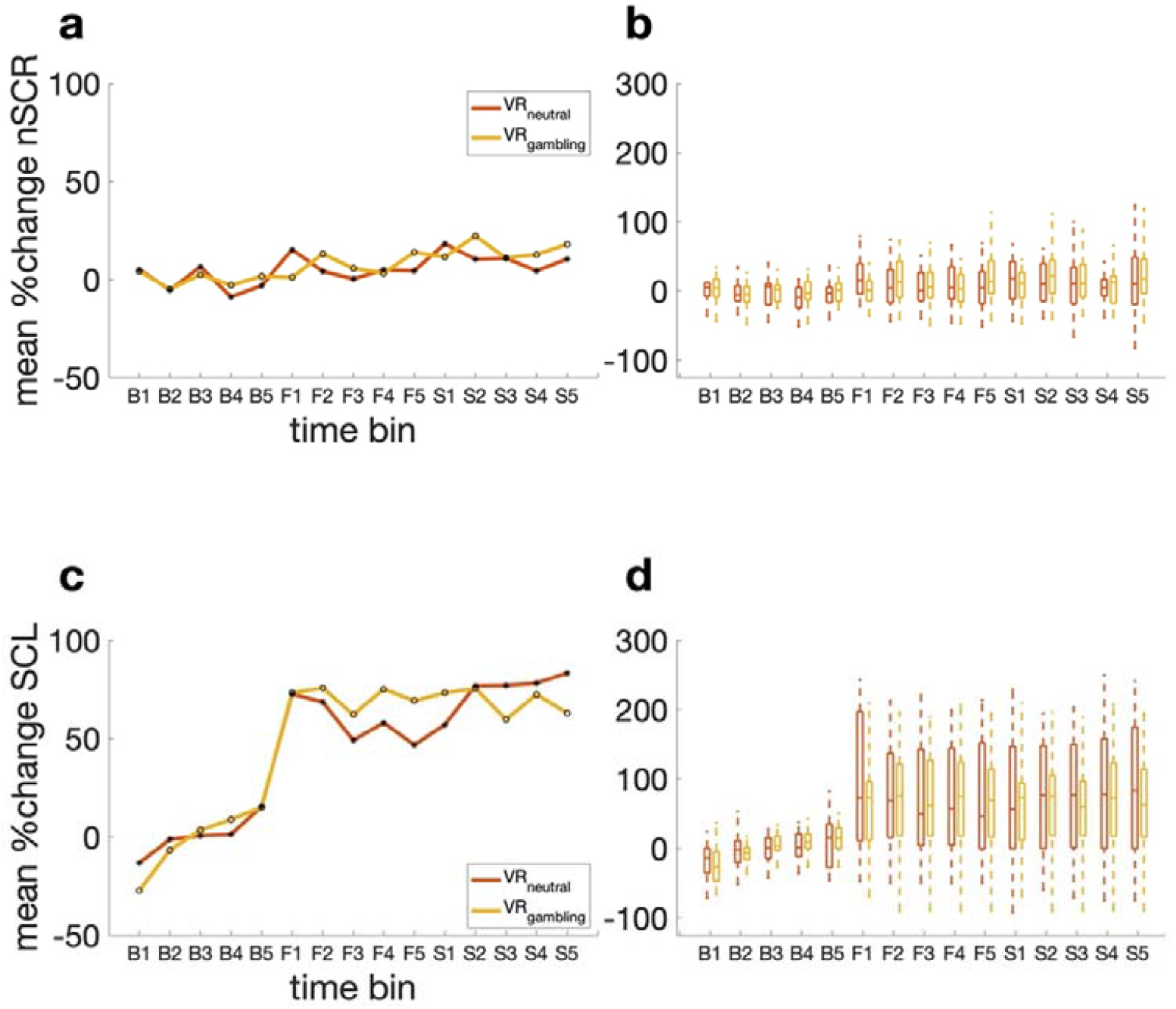
Results of the EDA measurements divided into 15 time points over the course of the baseline phase, measured before participants entered the VR-environments, and the first and second exploration phases. Each of the three phases is divided into five one-minute bins (B1-5: pre-VR baseline, F1-5: first exploration phase in VR, S1-5: second exploration phase VR). a: Median percent change from baseline mean for no. of spontaneous SCRs over all participants. b: Boxplot of percentage change from baseline mean for no. spontaneous SCRs over all participants. c: Median percent change from baseline mean of SCL over all participants. d: Boxplots of percentage change from base line mean of SCL over all participants.

## Discussion

Here we carried out an extensive investigation into the reliability of temporal discounting measures obtained in different virtual reality environments as well as standard lab-based testing. This design allowed us the joint assessment of physiological arousal and decision-making, an approach with potential applications to cue-reactivity studies in substance use disorders or behavioral addictions such as gambling disorder. Participants performed a temporal discounting task within two different VR-environments (a café environment and a casino/sports betting environment: VR_neutral_ vs. VR_gambling_) as well as in a standard computer-based lab testing session. Exposure to VR generally increased sympathetic arousal as assessed via electrodermal activity (EDA), but these effects were not differentially modulated by the different VR environments. Results revealed good to excellent test-retest reliability of model-based (*log(k)*) and model-free (AUC) measures of temporal discounting across all testing environments. However, the DDM_S_ parameters modelling latent decision processes showed substantially lower test-retest reliabilities between the three sessions. The split-half reliability within each session was mostly good to excellent indicating that the lower test-retest reliability was likely caused by the participants current state and not by factors within the modelling process itself.

To test how well temporal discounting, as a measure of choice impulsivity, performs in virtual environments we implemented a VR-design that is built for possible future application in a cue-reactivity context. Healthy controls displayed little evidence for systematic differences in choice preferences between the Lab-session and the VR-sessions. This was observed for model-free measures (AUC), as well as the *log(k)* parameter of the hyperbolic discounting model with the softmax choice rule and the drift diffusion model with non-linear drift rate scaling (DDM_S_). Model comparison revealed that the DDM_S_ accounted for the data best, confirming previous findings^[43,45,46,48]^. Although generally, discount rates assessed in the three sessions were of similar magnitude, in the DDM_S_ there was moderate evidence for reduced discounting (i.e., smaller values of *log(k)*) in the VR_neutral_ session. The reasons for this could be manifold. One possibility is that environmental novelty plays a role, such that perceived novelty of the VR_neutral_ session might have been lower than for the VR_gambling_ and Lab-sessions. Exposure to novelty can stimulated dopamine release^[77]^, which is known to impact temporal discounting^[78]^. Nonetheless, effect sizes were medium (.37 and .29) and the dBFs revealed only moderate evidence. Numerically, the mean *log(k)*’s of the softmax model showed the same tendency, but here effects were less pronounced. One possibility is that the inclusion of additional latent variables in the DDM_S_ might have ncreased sensitivity to detect this effect. There was also evidence for a session effect on the scaling parameter (*v*_coeff_). Here, the impact of trial-wise value differences on the drift rate was attenuated in the Lab-session, with dBFs revealing strong (VR_neutral_) or moderate evidence (VR_gambling_) for a reduction in *v*_coeff_ in the Lab-session. Again, effect sizes were medium. Nevertheless, the data suggest increased sensitivity to value differences in VR. This effect might be due to the option presentation in the Lab-session compared to the VR-sessions. The presentation of options within VR might have been somewhat more salient, which might have increased attention allocated to the value differences within the VR-sessions. However, this remains speculative until further research reproduces and further assesses these specific effects on the DDM parameters. Boundary separation (α), drift rate asymptote (*v*_*max*_) and starting point (*z*) showed little evidence for systematic differences between sessions. The only DDM_S_ parameter showing extreme evidence for a systematic difference between the lab- and VR-sessions was the non-decision time (r). This effect is unsurprising, as it describes RT components attributable to perception and/or motor execution. Given that indicating a response with a controller in three-dimensional space takes longer than a simple button press, this leads to substantial increases in τ during VR testing. Finally, the good test-retest reliability of *log(k)* from the DDM_S_ furthermore indicates that RTs obtained in VR can meaningfully be modeled using the DDM. The potential utility of this modeling approach in the context of gambling disorder is illustrated by a recent study that reported reduced boundary separation (a) in participants suffering from gambling disorder compared to healthy controls in a reinforcement learning task^[48]^. Given that there are mixed results when it comes to the effect of addiction related cues on RTs^[79–81]^, the effects of these cues on the latent decision variables included in the DDM could provide additional insights. Taken together, these results show that VR immersion in general does not influence participants inter-temporal preferences in a systematic fashion and might open up a road to more ecologically valid lab experiments, e.g., focusing on behavioral cue-reactivity in addiction. This is in line with other results showing the superiority of VR compared to classical laboratory experiments^[6]^.

The present data add to the discussion concerning the reliability of behavioral tasks^[9,50–53,55]^ in particular in the context of computational psychiatry^[15,82]^. To examine test-retest reliability, the three sessions were performed on different days and with a mean interval of 3.85 days between sessions. The test-retest reliability for the AUC and the *log(k)* parameter of the hyperbolic discounting model with softmax choice rule were both excellent. For the *log(k)* of the DDM_S_ the ICC was good, but slightly lower than for AUC and softmax. Nevertheless, the discount rate *log(k)* was overall stable regardless of the analytical approach. The ICC of .7 observed for the DDM_S_ was comparable to earlier studies on temporal discounting reliability^[52,53]^. Kirby and colleagues^[52]^ for instance demonstrated a reliability of .77 for a five-week interval and .71 for one year. This shows that at least over shorter periods from days to weeks, temporal discounting performed in VR has a reliability comparable to standard lab-based testing. Enkavi and colleagues^[49]^ stress that in particular difference scores between conditions (e.g. Stroop, Go-NoGo etc.), show unsatisfactory reliability due to the low between participants variation created by commonly used behavioral tasks. Assessment of difference scores was not applicable in the present study. Nevertheless, there was no positive evidence for systematic effects on *log(k)* (with the exception of the potential novelty effects discussed above), and the test-retest reliability between all conditions was at least good across analysis schemes, indicating short-term stability of temporal discounting measured in VR. It is worth noting, however, that temporal discounting shares some similarities with questionnaire-based measures. As in questionnaires, in temporal discounting tasks participants are explicitly instructed to indicate their preferences. This might be one reason why the reliability of temporal discounting is often substantially higher than that of other behavioral tasks^[49,52,53,55]^. Other parameters of the DDM_S_ showed lower levels of test-retest reliability. Especially the v_coeff_ parameters were less reliable, at least when estimated jointly with v_max_. In the DDM_L_, which does not suffer from potential trade-offs between these different drift rate components, the ICC of *v*_coeff_ was good (Supplementary Table S2). Similarly, here *log(k)* also showed an excellent ICC.

The substantially lower test-retest reliability exhibited by the parameters of the DDM_S_ that represent latent decisions processes, compared to *log(k)* or AUC warrants further discussion. Prior publications from our lab ^[24,41]^ have extensively reported parameter recovery of the DDMs model and revealed a good recovery performance. The low test-retest reliability is therefore unlikely to be due to poor identifiability of model parameters. One possible reason for this discrepancy between log(k)/AUC and the other parameters is that the tendency to discount value over time might be a stable trait-like factor, while the latent decision processes reflected in the other DDM_S_ parameters might be more substantially influenced by state effects. While this could explain the low test-retest reliability, it would predict that these parameters should nonetheless be stable within sessions. We addressed this issue in a further analysis of within-session split-half reliability (see Supplementary Tables S3-5). The results showed a good-to-excellent within-session stability for most parameters, with the drift rate coefficient *v*_coeff_ being a notable exception. This is compatible with the idea that latent decision processes reflected in the DDM_S_ parameters might be affected by factors that differ across testing days, but are largely stable within sessions, such as mood, fatigue or motivation.

VR has previously been used to study cue-reactivity in participants suffering from gambling disorder^[2,3,83]^, but also in participants experiencing nicotine^[84]^ and alcohol^[1]^ use disorders. Our experimental set-up extends these previous approaches in several ways. First, we included both a neutral and a gambling-related environment. This allows us to disentangle general VR effects from specific contextual effects. Second, our reliability checks for temporal discounting show that model-based constructs with clinical relevance for addiction^[18,23]^ can be reliably assessed when behavioral testing is implemented directly in the VR environment. Together, these advances might yield additional insights into the mechanisms underlying cue-reactivity in addiction, and contextual effects in psychiatric disorders more generally.

Understanding how addictions manifest on a computational and physiological level is important to further the understanding the mechanisms underlying maladaptive decision-making. Although alterations in neural reward circuits, in particular in ventral striatum and ventromedial prefrontal cortex, are frequently observed in gambling disorder, there is considerable heterogeneity in the directionality of these effects^[85]^. Gambling-related visual cues interfere with striatal valuation signals in participants suffering from gambling disorder, and might thereby increase temporal discounting^[12]^. In the present work, assessment of physiological reactivity to VR was limited to electrodermal activity (EDA). EDA is an index of autonomic sympathetic arousal, which is in turn related to the emotional response to addiction related cues^[39,86–88]^. The skin conductance level (SCL) is increased in participants with substance use disorders in response to drug related cues^[86]^. Additionally, it has been shown that addiction related cues in VR can elicit SCR responses in teen^[87]^ and adult^[88]^ participants suffering from a nicotine addiction. In our study, we mainly used this physiological marker to assess how healthy participants react to VR exposure. For the number of spontaneous responses in the EDA signal (nSCRs), the increase upon exposure to VR (B5 vs F1) was only significant in the VR_neutral_ environment. The effect size for the difference between both environments was medium. Given that the two starting areas of the VR-environments were identical, this difference might have been caused by random fluctuations. However, an increase in the number of spontaneous SCRs during VR immersion has been reported previously^[5]^ and thus warrants further investigation. The SCL, on the other hand, increased substantially upon exposure to VR, as indicated by a significant increase between the last minute of baseline recording (B5) and the first minute of the first exploration phase (F1). The effect sizes indicated a large effect. SCL then remained elevated throughout both exploration phases (F1 to S5) but did not increase further when the virtual café/casino area was entered. These results suggest that exposure to VR increases sympathetic arousal as measured with SCL in healthy control participants independent of the presented VR environment.

There are several limitations that need to be acknowledged. First, there was considerable variability in test-retest intervals across participants. While most of the sessions were conducted within a week, in some participants this interval was up to three weeks, reducing the precision of conclusions regarding temporal stability of discounting in VR. Other studies, however, have used intervals ranging from five to fifty-seven weeks^[52]^ or three months^[53]^, and have reported comparable reliabilities. Moreover, there is evidence for a heritability of temporal discounting of around 30 and 50 percent at the ages of 12 and 14 years respectively^[89]^. This increases the confidence in the results obtained here. Nevertheless, a more systematic assessment of how long these trait indicators remain stable in VR would be desirable and could be addressed by future research. Second, the sample size was lower compared to larger studies conducted online^[49]^, and the majority of participants was female. Both factors limit the generalizability of our results. However, large-scale online studies have shortcomings of their own, including test batteries that take multiple hours and/or multiple sessions to complete^[49,50]^, potentially increasing participants’ fatigue, and which might have detrimental effects on data quality. We also note that the present sample size was sufficiently large to reveal stable parameter estimates, showing that in our design participants performed the task adequately. Thirdly, the immersion in VR might have been reduced by the available physical lab space. To ensure safety, the experimenter had to at times instruct participants to stay within the designated VR-zone. This distraction might have reduced the effects caused by the VR-environments, because participants were not able to fully ignore the actual physical surroundings. Additionally, it might have influenced the EDA measurements in an unpredictable way. Future research would benefit from the implementation of markers within the VR-environments in order to ensure safety without breaking immersion. Moreover, participants had to spend about thirty minutes in the full VR-setup. The behavioral tasks were presented after the exploration phase, such that participants might have been fatigued or experienced discomfort during task completion. Finally, the study at hand did not include participants that gamble frequently or are suffering from gambling disorder and is therefore not a cue-reactivity study itself, but rather a methodological validation for future studies using this and similar designs. Due to the fact that participants here were supposed to be fairly unfamiliar with gambling environments this study could not determine how ecologically valid the gambling environment actually is. This needs to be addressed in future research. In relation to that, cue-reactivity in gambling disorder is determined by many individual factors^[37]^. The VR-design presented here is designed for slot machine and sports betting players, and thus not applicable for other forms of gambling.

Overall, our results demonstrate the methodological feasibility of a VR-based approach to behavioral and physiological testing in VR with potential applications to cue-reactivity in addiction. Healthy non-gambling control participants showed little systematic behavioral and physiological effects of the two VR environments. Moreover, our data show that temporal discounting is reliable behavioral marker, even if tested in very different experimental settings (e.g. standard lab testing vs. VR). It remains to be seen if such gambling-related environments produce cue-reactivity in participants suffering from gambling disorder. However, results from similar applications have been encouraging^[2,3]^. These results show the promise of VR applications jointly assessing of behavioral and physiological cue-reactivity in addiction science.

## Acknowledgments

We thank Mohsen Shaverdy and Diego Saldivar for the implementation of the VR environments and task programming and all members of the Peters Lab at the University of Cologne for helpful discussions. This work was supported by Deutsche Forschungsgemeinschaft (DFG, grant PE1627/5-1 to J.P.).

## Additional Information

The authors declare no competing interests.

## Author Contributions

Conceptualization: J. P., L. B.

Data curation: L. B.

Formal analysis: L. B.

Funding acquisition: J. P.

Investigation: L. B., L. S.

Methodology: J. P., L. B.

Project administration: L. B.

Writing – original draft: L. B.

Writing – review & editing: J. P., L. B., L. S.

## Supplementary Materials

**Supplementary Figure S1.**
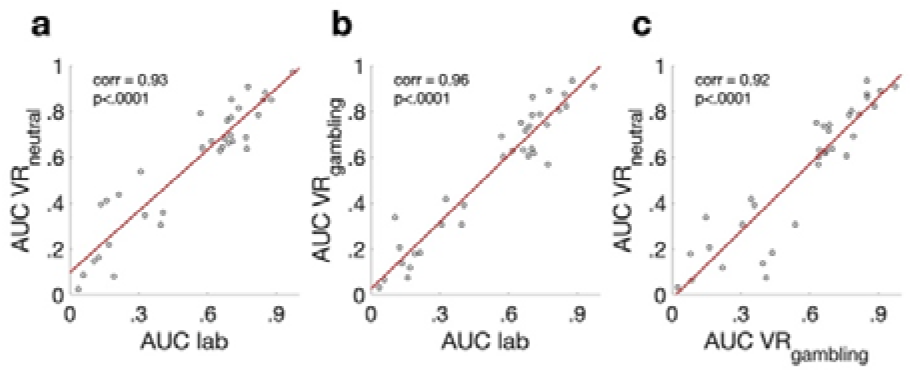
Scatterplots of the individual participants AUC values. a) lab vs VR_neutral_ b) lab vs VR_gambling_ c) VR_gambling_ vs VR_neutral_.

**Supplementary Figure S2.**
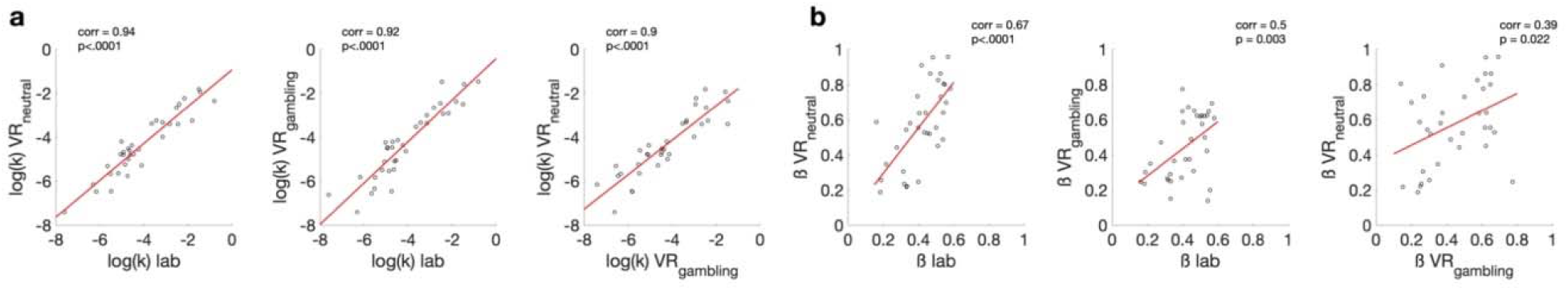
Scatterplots of the mean of the individual participants parameter posterior distributions for the parameters of the hyperbolic discounting model with the softmax choice rule (see equation 1 and 2). a) *log(k)* b) softmax ß.

**Supplementary Figure S3.**
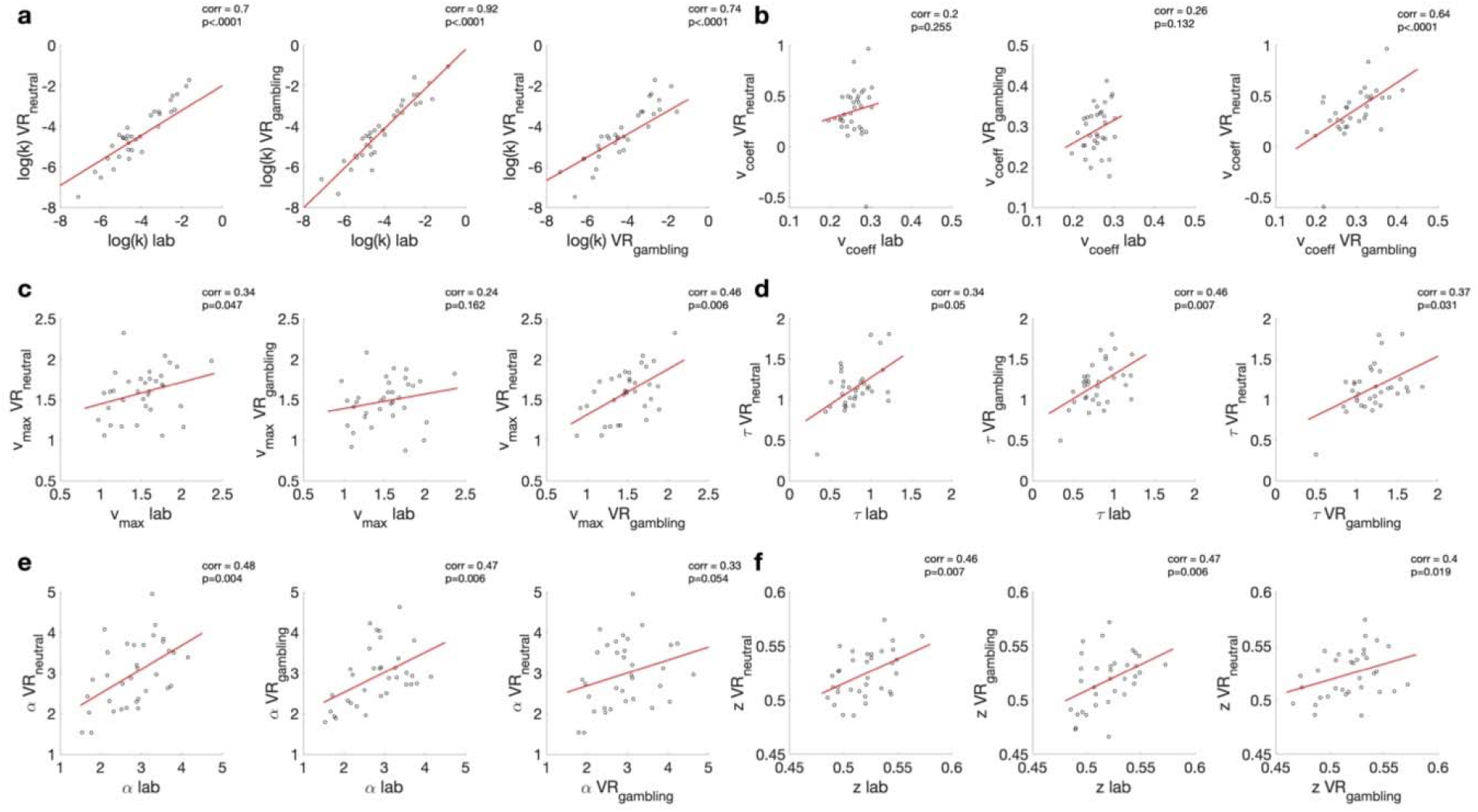
Scatterplots of the mean of the individual participants parameter posterior distributions for the parameters of the DDM_S_ temporal discounting model. a) *log(k)* b) *v*_coeff_ c) *v*_max_ d) tau e) a f) z.

**Supplementary Table S1.**
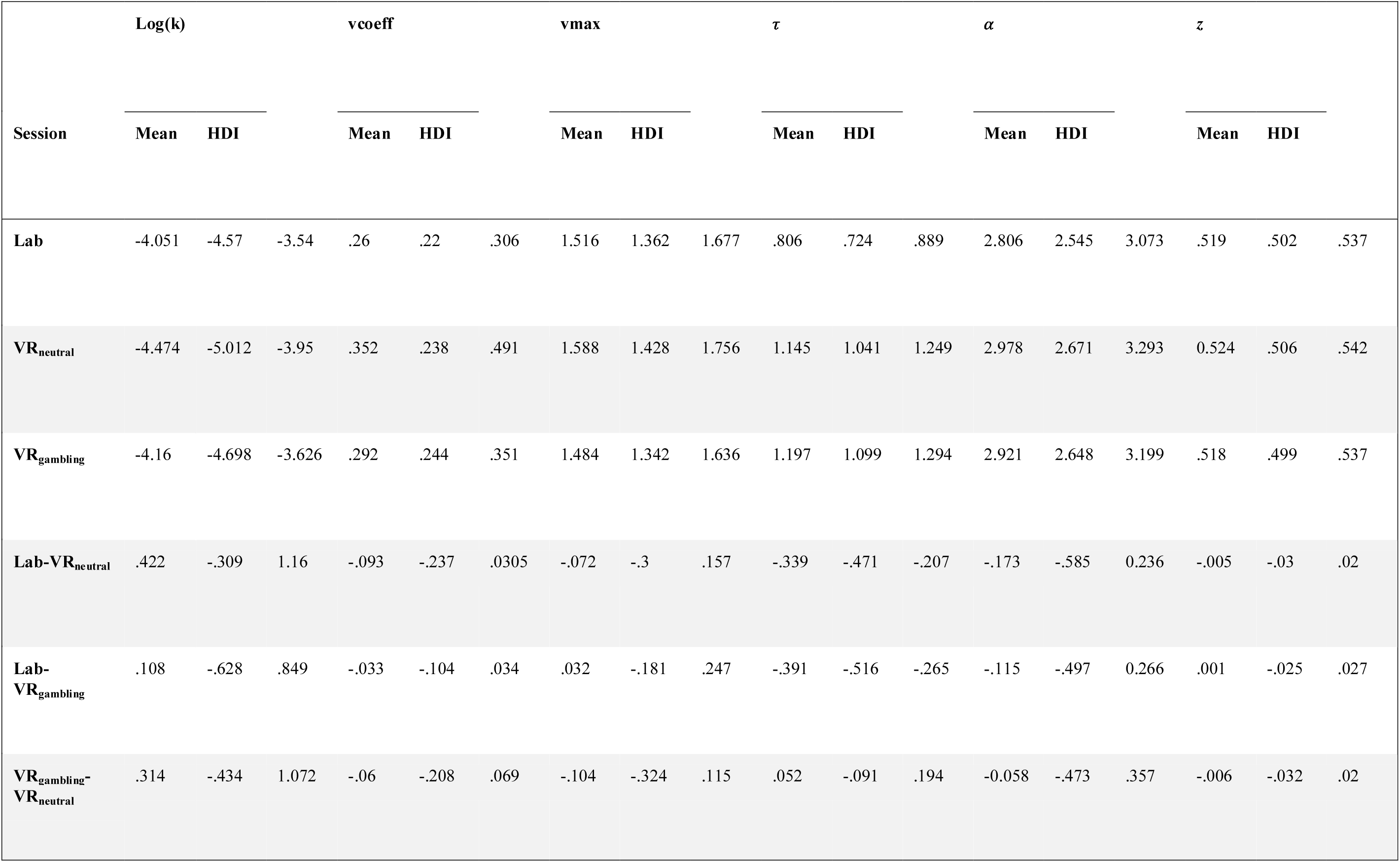
Means and 95% HDIs of the posteriors of the parameters from the DDM_s_ model.

**Supplementary Table S2.** Summary of the results of the ICC analysis of the DDM_L_ parameters.

| Parameter | ICC | p | Lower bound | Upper Bound |
| --- | --- | --- | --- | --- |
| **log(k)** | .92 | <.001 | .88 | .95 |
| **$\nu_{\text{coeff}}$** | .65 | <.001 | .5 | .77 |
| **$\tau$** | .15 | .067 | -.014 | .35 |
| **$\alpha$** | .36 | <.001 | .19 | .55 |
| **$z$** | .60 | <.001 | .45 | .74 |

**Supplementary Table S3.** Summary of the results of the split-half ICC analysis of the DDM_S_ parameters within the lab-session.

| Parameter | ICC | p | Lower bound | Upper Bound |
| --- | --- | --- | --- | --- |
| **log(k)** | .97 | <.001 | .96 | .97 |
| **$\nu_{\text{coeff}}$** | .25 | .069 | -.029 | .5 |
| **$\nu_{\text{max}}$** | .76 | <.001 | .61 | .86 |
| **$\tau$** | .92 | <.001 | .86 | .95 |
| **$\alpha$** | .94 | <.001 | .9 | .97 |
| **$z$** | .48 | .002 | .23 | .67 |

**Supplementary Table S4.** Summary of the results of the split-half ICC analysis of the DDM_S_ parameters within the VR_neutral_-session.

| Parameter | ICC | p | Lower bound | Upper Bound |
| --- | --- | --- | --- | --- |
| **log(k)** | .73 | <.001 | .57 | .84 |
| **$\nu_{\text{coeff}}$** | .005 | .49 | -.092 | .29 |
| **$\nu_{\text{max}}$** | .65 | <.001 | .46 | .79 |
| **$\tau$** | .94 | <.001 | .9 | .97 |
| **$\alpha$** | .9 | <.001 | .82 | .94 |
| **$z$** | .25 | .075 | -.036 | .49 |

**Supplementary Table S5.** Summary of the results of the split-half ICC analysis of the DDM_S_ parameters for the parameters within the VR_gambling_-session.

| Parameter | ICC | p | Lower bound | Upper Bound |
| --- | --- | --- | --- | --- |
| **log(k)** | .96 | <.001 | .56 | .8 |
| **$v_{\text{coeff}}$** | -.1 | .73 | -.053 | .3 |
| **$v_{\text{max}}$** | .58 | <.001 | .16 | .52 |
| **$\tau$** | .94 | <.001 | .019 | .38 |
| **$\alpha$** | .92 | <.001 | .24 | .59 |
| **$z$** | .36 | .016 | .22 | .58 |

